# Validation of Extracellular Vesicles in FACTORFIVE’s EXOFECTA LyoBeads

**DOI:** 10.1101/2025.11.17.688893

**Authors:** David L Stachura, Kristen F Rueb, Dustin Hadley, Batuhan Bozkurt, Randy P Carney, John Aylworth

## Abstract

**Objective:** The objective of this project was to determine if extracellular vesicles (EVs) were present in FACTORFIVE’s EXOFECTA LyoBeads to validate flash freezing and subsequent lyophilization as an effective EV preservation method.

**Summary:** EVs were isolated from resuspended EXOFECTA LyoBeads using differential ultracentrifugation and subjected to further characterization techniques such as nanoparticle tracking analysis (NTA), single particle flow cytometry, and cryo-electron microscopy (cryo-EM). First, NTA revealed particles with size distributions and concentrations consistent with the expected characteristics of EVs, indicating successful isolation from samples. Secondly, flow cytometry was performed using antibodies targeting the canonical EV tetraspanins CD9, CD63, and CD81. Detection of these markers supported the presence of enriched EVs in the samples. Finally, cryo-EM was performed to confirm the existence of vesicular bodies and their morphology. Visualization of the samples revealed membrane-enclosed particles with a visible lipid bilayer, consistent with EVs. Overall, the results support EVs were in EXOFECTA LyoBeads, indicating EV retention in the final product after FACTORFIVE’s method of flash freezing, stabilizing, and lyophilization.

## Background

Extracellular vesicles (EVs) are membrane-bound particles naturally released from cells, which cannot replicate and play a role in intercellular communication. They have been identified from almost every cell or organism examined, indicating their ubiquity in all empires of living things (reviewed in Woith et al., 2019). EVs are commonly referred to as “exosomes”, but exosomes are only a subset of EVs that derive from a specific biogenic origin whereby the multivesicular body in the cell fuses with the plasma membrane. In addition to exosomes, cells also secrete numerous different EV subtypes (Buzas, 2022), with more and more diversity emerging regularly as the field advances. Since exosome-type EVs can only be identified by visualizing their biogenesis in cells, most assays used in the laboratory to examine exosomes are actually examining heterogeneous EVs. Due to this fact, the International Society for Extracellular Vesicles (ISEV) recommends to refer more generally to these particles simply as EVs (as we do in this report) or even extracellular particles (EPs) (Welsh et al., 2024).

EVs have attracted significant interest due to their ability to carry bioactive molecules. This capability has made them promising candidates for diagnostic, prognostic, and therapeutic applications (see van Niel et al., 2018; Simons and Raposo, 2009). In essence, EVs are discrete packages that can contain cellular proteins, glycans, signaling molecules, growth factors, DNA/RNA, including microRNAs (which are important for regulating gene expression). They are wrapped in a lipid bilayer, so they can effectively fuse with neighboring or distant cells and release their cargo directly or potentiate surface-mediated signal transduction via binding. EVs enable long distance communication, transmitting their cargo with minimal diffusion, dilution, or degradation compared to unpackaged cargo (Wolf and Casadevall, 2014). Some EVs also contain matrix remodeling enzymes on their surface such as matrix metalloproteinases and hyaluronidases (Nawaz et al., 2018). These enzymes play roles in extracellular matrix (ECM) turnover and composition in several physiological and pathological conditions. Additionally, EVs play active roles in tissue regeneration by stimulating repair processes (Fatima et al., 2017).

Mesenchymal stem cells (MSCs) are adult stem cells that can regenerate more of themselves and differentiate into adipocytes (fat), osteocytes (bone), and chondrocytes (cartilage) (Uccelli et al., 2008). They can also differentiate into fibroblasts, which are important components of connective tissue (X. Tanaka et al., 2022). MSCs were first discovered in the bone marrow, but are also present in the hypodermis, which is the lowest, fat-filled layer of skin; these are called adipose-derived MSCs (ADMSCs) (Zuk et al., 2001). There is a high frequency of ADMSCs in fat (Strioga et al., 2012 and Hass et al., 2011), and over 1.5 million liposuction surgeries are performed globally every year; the fat removed is regarded as waste and can be collected without additional risk to the donor. ADMSCs can differentiate into epidermal stem cells (Derby et al., 2014) and fibroblasts (X. Tanaka et al., 2022), indicating that they are likely the most important stem cell population for skin.

ADMSCs communicate to surrounding cells by secreting signaling proteins, as well as ECM molecules. ADMSCs contribute to the regeneration of skin (Lynch and Pei, 2014; Kim et al., 2008; Zhang et al., 2014) and secrete ECM proteins like collagen and elastin that reduce fine lines and wrinkles (Novoseletskaya et al., 2020), as well as signaling molecules that instruct cells to grow and migrate to an injury (Pittenger et al., 2019). Proteins produced by ADMSCs remodel scar tissue, increase blood flow, and modulate the immune system (Melief et al., 2013). Additionally, they secrete antioxidants (Kim et al., 2008 and Zhang et al., 2014). Transplantation studies indicate young ADMSCs rejuvenate aging skin (Bernardini et al., 2015 and Gennai et al., 2017) and secrete growth factors that promote cellular homeostasis and repair (Park et al., 2008). Studies show that using the proteins secreted from ADMSCs helps with a multitude of skin issues, including reducing the appearance of aging. Importantly, ADMSCs also secrete EVs that activate signaling pathways involved in wound healing (Zhou et al., 2021), encourage angiogenesis (Liang et al., 2016), and mediate immune responses (Matwiejuk et al., 2025). In addition, EVs induce the expression of several essential growth factors in ADMSCs (reviewed in Fang et al., 2019). For these reasons, cellular secretions from ADMSCs hold great promise to accelerate skin wounding and regeneration and increase skin health.

Unfortunately, EVs are notoriously finicky biomolecular assemblies. They are sensitive to temperature and are typically stored below −80°C for long-term preservation (reviewed in Ahmadian et al., 2024). Studies indicate that rapid freezing in liquid nitrogen preserves EV particle numbers compared to slow freezing (Wright et al, 2022) and that lyophilization maintains EV size, morphology, protein content, and bioactivity when accompanied by appropriate cryoprotectants like trehalose (Lyu et al., 2022 and Neupane et al., 2021). Based on these previous studies, we hypothesized that it was possible to make a stabilized cosmetic product that contained EVs.

We obtained consent to isolate ADMSCs from human lipoaspirate samples from patients between 20 and 25 years of age and then tested those cells to ensure they had no communicable diseases. Then, ADMSCs were grown with a patent-pending process to generate Dynamic Culture Medium (DCM). In essence, DCM contains secreted proteins including growth factors, cytokines, immunomodulatory proteins, antioxidants, ECM proteins, and ECM remodeling proteins. DCM also contains EVs.

To generate a product that captured all the proteins and EVs from ADMSC secretions, we created EXOFECTA. Simply, EXOFECTA is DCM that has added trehalose to stabilize EVs, as well as GHK-Cu copper peptides which are proven to speed wound healing, activate stem cells, increase antioxidant activity, repair fibroblasts, increase collagen production, and rejuvenate skin (reviewed in Pickart and Margolina, 2012). This solution is flash frozen in liquid nitrogen, creating small 20 µL droplets. These droplets are then lyophilized, creating EXOFECTA LyoBeads. After freeze drying, the LyoBeads are packed in sterile glass vials. When needed, they are reconstituted in sterile, injection-grade hyaluronic acid and utilized for skincare treatments.

In this product note, we report a simple characterization analysis in collaboration with the UC Davis Extracellular Vesicles Acquisition and Analysis Core (EV Core). We utilized the Minimal Information for Studies of Extracellular Vesicles (MISEV) 2023 guidelines to design and perform these experiments. MISEV guidelines focus on characterizing EVs using multiple complementary platforms to assess physical characteristics, biochemical composition, and mode of biogenesis (e.g., if specific subpopulation nomenclature such as exosomes or microvesicles is desired). In alignment with the MISEV guidelines, this report emphasizes methodological transparency and standardization. Adhering to these community-established criteria enhances data reproducibility, facilitates comparison across studies, and supports the credibility of findings for regulatory and translational purposes. This study was completed in partial compliance of core MISEV 2023 criteria, with further orthogonal and functional validations to follow. Overall, we report that the characterization results are consistent with those expected for enriched EVs, supporting their presence in EXOFECTA.

## Methods

### Samples

The samples were EXOFECTA LyoBeads provided to the EV Core by FACTORFIVE.

### EV Isolation

Samples underwent differential ultracentrifugation (UC) to enrich EVs (**Figure 1**). The process included the following steps:

1. *Initial clarification and removal of cells, debris, and larger vesicles (e.g., microvesicles):* Samples were centrifuged at 300 x g for 10 min, 2,000 x g for 15 min, and 10,000 x g for 30 min. Each time, the supernatant was carried over to the next step.
2. *EV pelleting*: Supernatant from the previous step was centrifuged at 120,000 x g for 70 min in a Beckman Coulter Optima TLX. The supernatant was collected and retained.
3. *EV washing*: The pellet from the previous step was resuspended with filtered phosphate buffered saline (PBS) and centrifuged again at 120,000 x g for 70 min. The supernatant was collected and retained. The pellet was resuspended, aliquoted, and frozen at −80°C until used for downstream characterization.

**Figure 1:**
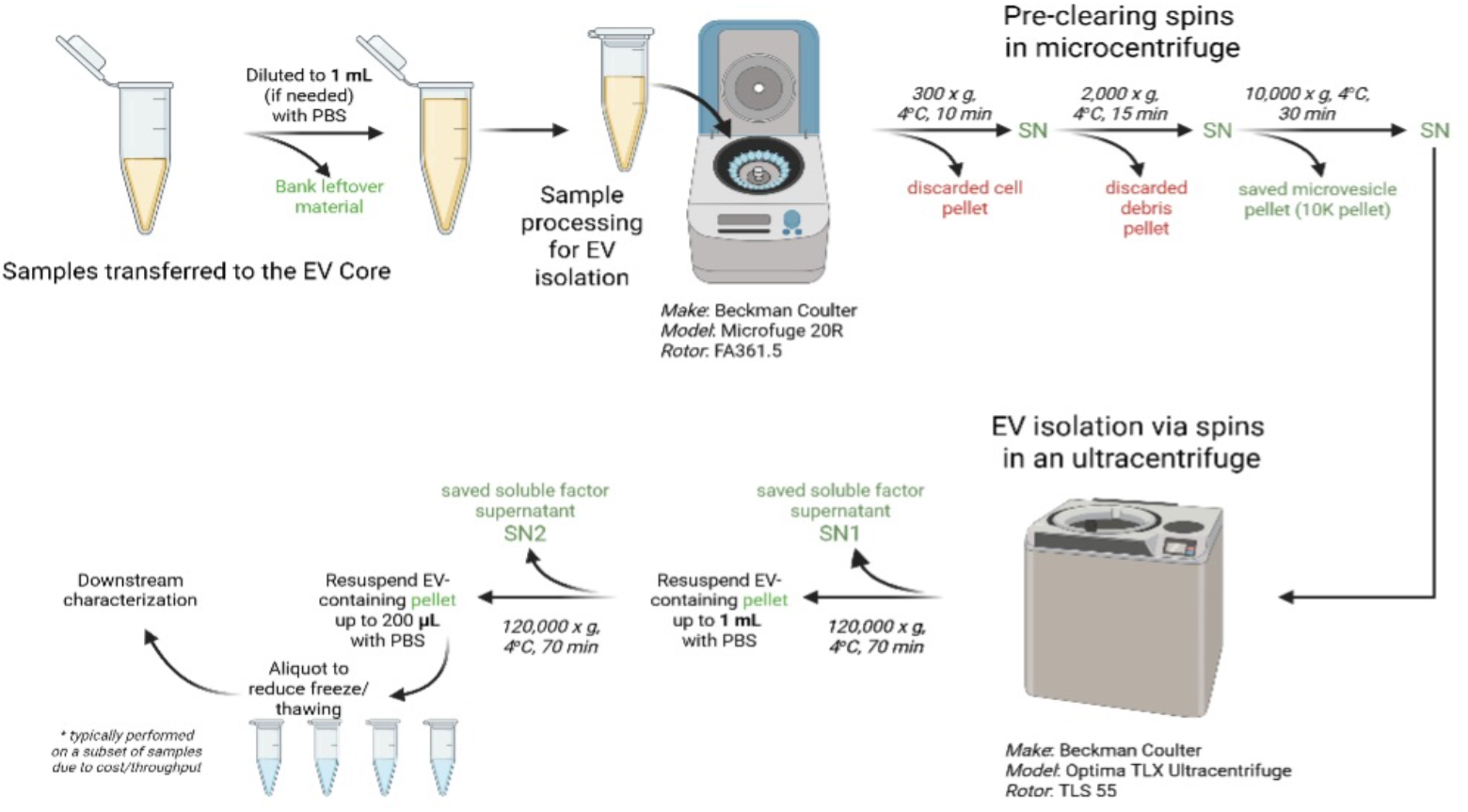
Experimental design for EV isolation and validation

### Nanoparticle Tracking Analysis (NTA)

Samples were analyzed using a Malvern NanoSight LM14. Prior to each measurement, samples were diluted with 0.02 µm filtered PBS to fall within the optimal particle concentration range recommended by the manufacturer for reliable quantification. Measurements were performed using a camera level/laser power setting of 13 and a detection threshold of 2. Each sample was analyzed from three technical replicates, with each replicate consisting of a 90 sec video capture. Key metrics obtained included particle concentration (particles/mL), size distribution, and mode diameter.

### Flow Cytometry

Flow cytometry was performed using the Beckman Coulter CytoFLEX Nano, a nanoflow cytometer with sufficient sensitivity to detect and analyze single particles. This approach aligns with MISEV 2023 recommendations for the detection of EV surface markers. Samples were stained with a panel of antibodies targeting the canonical tetraspanin markers CD9, CD63, and CD81 (Andreu and Yanez-Mo, 2014).

Samples were incubated in the dark at room temperature for 90 min. Following incubation, excess antibodies were removed using Izon qEV 35 nm SEC columns. An antibody-only negative control and a HEK293T-derived EV positive control were included to support marker specificity and instrument QC.

Data were analyzed using fluorescence intensity thresholds and gating strategies informed by the negative control. The presence of CD9, CD63, and CD81 signals support the EV nature of the particles analyzed.

### Cryo-Electron Microscopy (Cryo-EM)

Cryo-EM imaging was performed to confirm the existence of vesicular bodies and their morphology. Samples were applied to glow-discharged grids and plunge frozen using the PELCO easiGlow system and the Vitrobok Mk IV. Grids were imaged using a 200 kV transmission electron microscope, the TFS Glacios Cryo-TEM, at magnifications ranging from 11,000x to 45,000x.

Images were evaluated for EV characteristics such as round, lipid bilayer-enclosed structures. Size and morphology were annotated and measured with ImageJ. Diameter used the outer leaflet to measure Feret diameter.

## Results

### Yield and Size Distribution of EVs with NTA

Two individual samples of EXOFECTA LyoBeads were processed and enriched for EVs (**Figure 1**) and examined with NTA (**Figure 2**). NTA indicated an average number of 1.10^10^ EVs that were 104.14 nm in size present in the two samples examined (**Figure 2** and **Table 1**). The yield and size of particles observed from NTA suggested these particles were EVs that should be examined further by flow cytometry and cryo-EM.

**Table 1:** Quantitation of EV numbers and size present in EXOFECTA LyoBead samples. Numbers are the average of sample B and C. Data shown in Figure 1.

| Sample | Avg. Conc. / SD (Particles/mL) | Avg. Mode Size / SD (nm) |
| --- | --- | --- |
| EXOFECTA LyoBeads | $1.10^{10} / 5.2^8$ | 104.14 / 4.67 |

**Figure 2:**
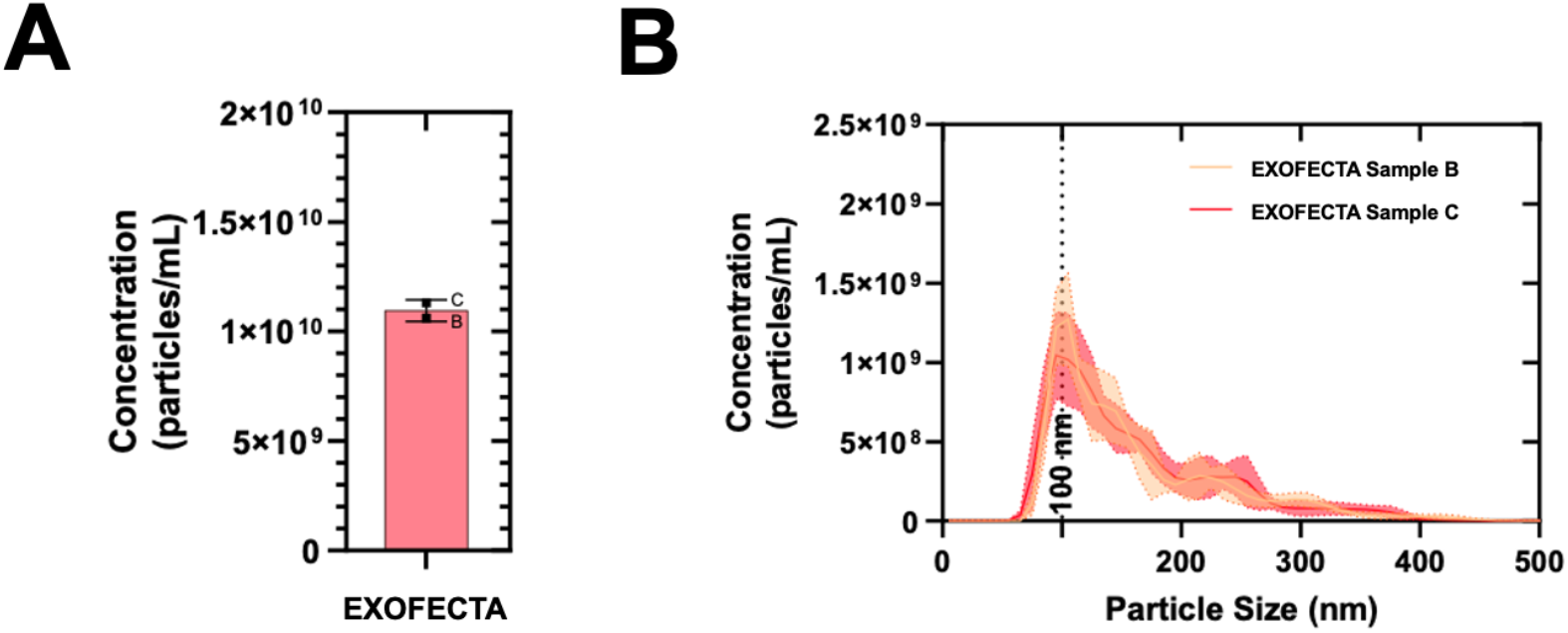
EVs are present in reconstituted EXOFECTA LyoBeads as measured by NTA. A) Average of EVs detected by NTA samples B and C denoted by the red bar, with SD indicated by the error bars. B) Particle size distribution of EXOFECTA sample B (orange line) and sample C (red line) indicate the presence of EVs in the samples.

### Expression of Tetraspanins on EVs with flow cytometr*y*

Flow cytometry was used to detect the presence of the canonical EV surface markers CD9, CD63, and CD81. All samples demonstrated detectable expressions of these tetraspanins, supporting the presence of EVs (**Figure 3**).

**Figure 3:**
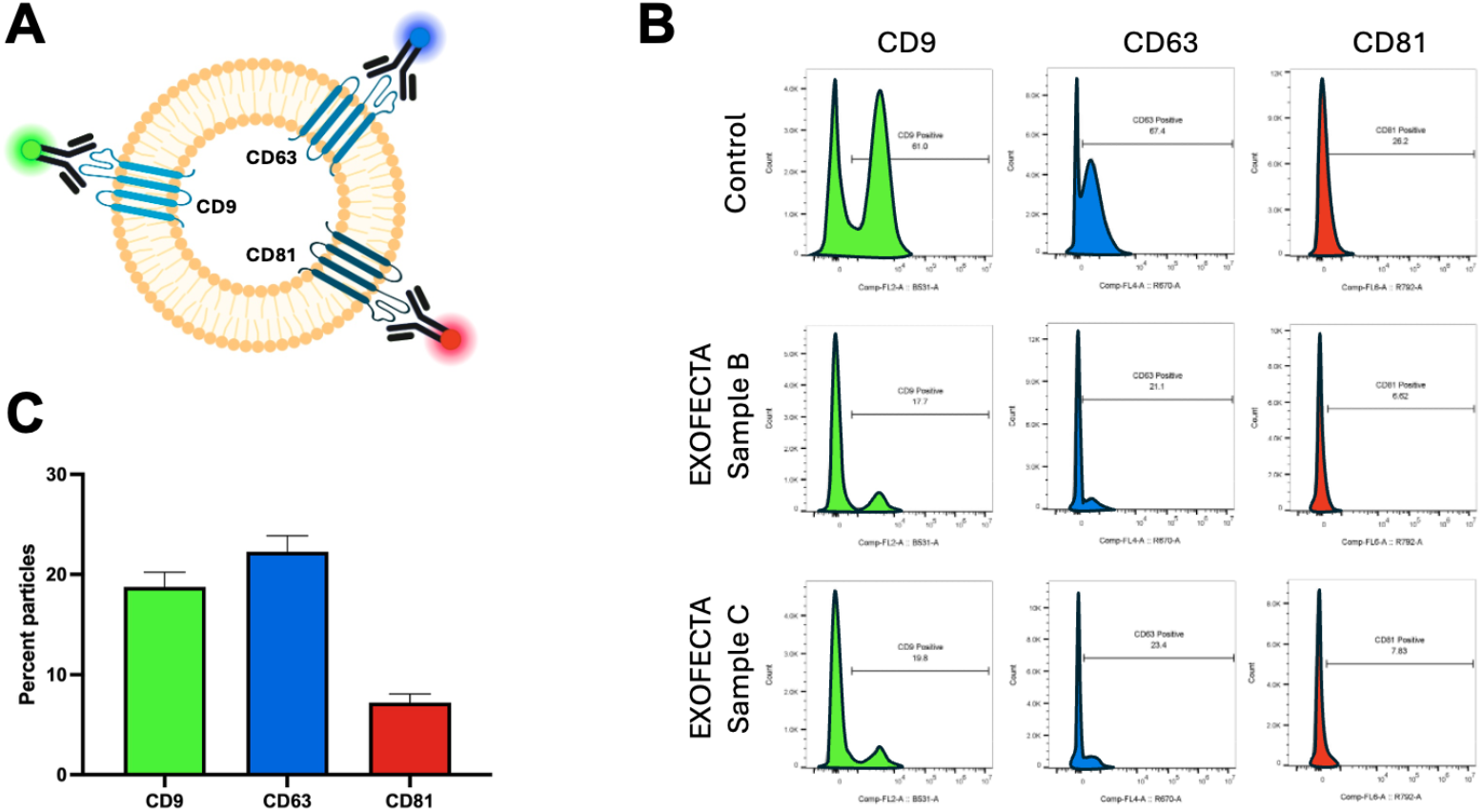
EVs are present in reconstituted EXOFECTA LyoBeads as measured by flow cytometry for CD9, CD63, and CD81. **(A)** Diagram showing the mode of measurement via primary-labeled fluorescent antibodies for EVs with CD9 (green), CD63 (blue), and CD81 (red) tetraspanins present. **(B)** Top row shows HEK293T-derived positive controls for CD9 (green, left column), CD63 (blue, middle column), and CD81 (red, right column) tetraspanins. EXOFECTA sample B (middle row) and EXOFECTA sample C (bottom row) show the presence of the three proteins on the surface of EVs. **(C)** Quantification of CD9 (green bars), CD63 (blue bars), and CD81 (red bars) with SD (error bars). Data are from counts in **(B)**.

### Morphology with cryo-EM analysis

To examine if EVs from EXOFECTA exhibited typical morphology of EVs, cryo-EM was performed. Vesicular structures consistent with EVs were observed in EXOFECTA samples post-EV enrichment (**Figure 4**).

**Figure 4:**
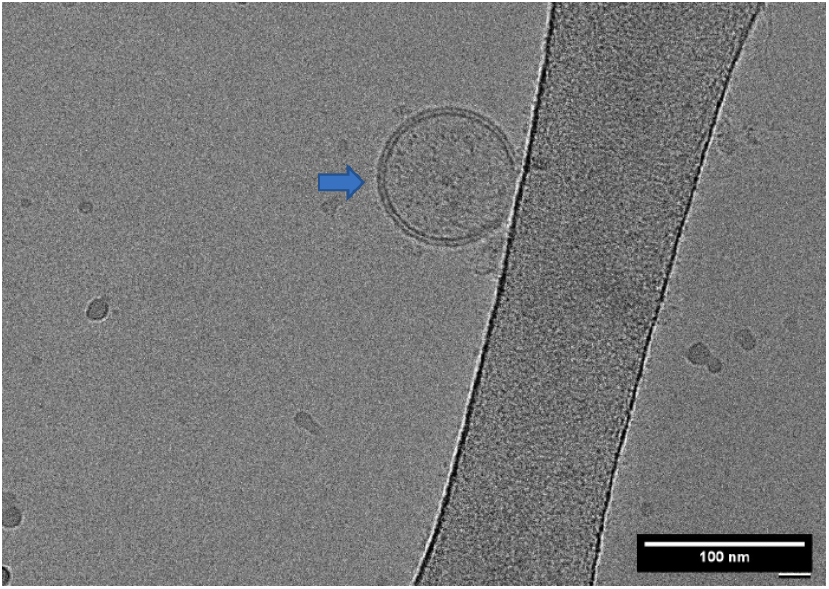
Unilamellar EV identified after cryo-EM analysis. Blue arrow indicates the presence of an EV on a cryo-EM grid.

The EV sizes observed during cryo-EM analysis were notably smaller than the 100 nm mode suggested by NTA results (**Figure 5**). These data indicated that there was a large fraction of EVs smaller than the limit of detection of the NTA device (compare to **Figure 2** and **Table 1**).

**Figure 5:**
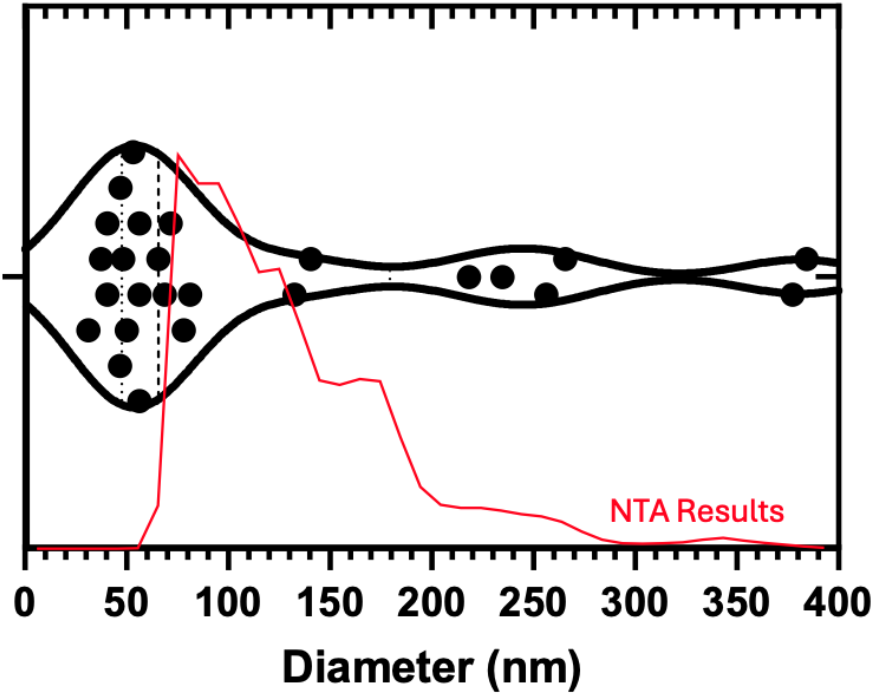
Many EVs are below the detection limit of NTA. 25 individual EVs were measured via cryo-EM (black dots and violin plot) from the lyophilized sample, plotted against results obtained from NTA (thin red line) from a frozen sample.

## Discussion

EVs were successfully identified in the lyophilized EXOFECTA LyoBead samples. NTA results showed similar particle concentrations between previously frozen and lyophilized samples (data not shown). Flow cytometry confirmed tetraspanins commonly observed with EVs were present in the samples. Cryo-electron microscopy of the samples revealed membrane-enclosed particles with a visible lipid bilayer, consistent with EVs. Additionally, measurement of EV size with cryo-EM indicated that NTA analysis is likely underestimating the size and number of EVs present in the samples.

The study meets several MISEV criteria, with further characterization recommended to fully satisfy these community-established guidelines. Of the characterization performed, NTA’s detection threshold of ∼70 nm underrepresents smaller EVs and their heterogeneity. Morphological analysis via cryo-EM can address some uncertainty for the presence of smaller EVs and the impact of lyophilization. Molecular analysis via flow cytometry corroborates evidence of EVs due to the presence of EV-associated proteins in the lipid bilayer. Future plans to characterize EXOFECTA LyoBeads will focus on protein and miRNA contents of the isolated EVs, as well as performing additional orthogonal measures of size, concentration, and molecular analysis.

